# A novel mutual information estimator to measure spike train correlations in a model thalamocortical network

**DOI:** 10.1101/289512

**Authors:** Ekaterina D. Gribkova, Baher A. Ibrahim, Daniel A. Llano

## Abstract

The impact of thalamic state on information transmission to the cortex remains poorly understood. This limitation exists due to the rich dynamics displayed by thalamocortical networks and because of inadequate tools to characterize those dynamics. Here, we introduce a novel estimator of mutual information and use it to determine the impact of a computational model of thalamic state on information transmission. Using several criteria, this novel estimator, which uses an adaptive partition, is shown to be superior to other mutual information estimators with uniform partitions when used to analyze simulated spike train data with different mean spike rates, as well as electrophysiological data from simultaneously recorded neurons. When applied to a thalamocortical model, the estimator revealed that thalamocortical cell T-type calcium current conductance influences mutual information between the input and output from this network. In particular, a T-type calcium current conductance of about 40 nS appears to produce maximal mutual information between the input to this network (conceptualized as afferent input to the thalamocortical cell) and the output of the network at the level of a layer 4 cortical neuron. Furthermore, at particular combinations of inputs to thalamocortical and thalamic reticular nucleus cells, thalamic cell bursting correlated strongly with recovery of mutual information between thalamic afferents and layer 4 neurons. These studies suggest that the novel mutual information estimator has advantages over previous estimators, and that thalamic reticular nucleus activity can enhance mutual information between thalamic afferents and thalamorecipient cells in the cortex.

## Introduction

Estimating the degree to which a signal is conserved as it passes through a neural network requires methods that are both sensitive to spike timing and that take into account nonlinear dependencies between spike trains. These requirements are particularly necessary in thalamocortical networks, which encode stimuli nonlinearly (Kayser et al. 2001; MacLean et al. 2005; Miller et al. 2014; Watson et al. 2008) and rely on the precise timing of inputs to process sensory information (Rose and Metherate 2005). Sensory thalamocortical (TC) neurons receive input from retina or caudal sensory structures and project to well-defined areas of the cerebral cortex. TC neurons display at least two firing modes: tonic and burst mode, and tonic mode has generally been regarded as a high-fidelity transmission state to relay sensory information to the cortex (Jones 2007; Kim and McCormick 1998; Llinás and Steriade 2006; McCormick and Feeser 1990; Sherman 2001; Steriade and Llinás 1988). TC cells are also strongly synaptically interconnected with the thalamic reticular nucleus (TRN), which comprises a shell of GABAergic neurons that partially surrounds the thalamus. The TRN has been implicated in a wide range of brain functions, including the production of sleep spindles, modulation of arousal and attention and, under pathological conditions, production of absence seizures (Destexhe et al. 1999; Destexhe et al. 1993; Halassa et al. 2014; Halassa et al. 2011; Huguenard 1998; McAlonan et al. 2008; McCormick and Contreras 2001). Traditional models of thalamic processing have postulated the presence of reciprocal connectivity between TC and TRN neurons, and that this connectivity forms the basis for oscillations such as spindles and spike-wave discharges in absence epilepsy (Destexhe et al. 1998; Huguenard 1998; Steriade et al. 1993). However, more recent data have revealed non-reciprocal connectivity, which may serve as a substrate to adjust the gain and filter properties of TC cells or to select particular groups of TC cells to meet cognitive demands (Crabtree and Isaac 2002; Kimura 2014; Pinault and Deschênes 1998; Willis et al. 2015; Zikopoulos and Barbas 2012). We recently explored a non-reciprocal (“open-loop”) model and found that total spike counts in the output of the model, a layer 4 (L4) cortical cell, were paradoxically enhanced by intermediate rates of TRN activity, and that this enhancement was dependent on both the actions of TRN neurons and nonlinear T-type calcium currents (T-currents) in TC cells (Willis et al. 2015), consistent with recent physiological findings (Whitmire et al. 2017; Whitmire et al. 2016). Given the impact of T-channel-mediated bursting behavior on enhancement of information transmission through a TC network (Reinagel et al. 1999), and the theoretical benefits of bursting on signal encoding (Denning and Reinagel 2005; Lisman 1997; Mukherjee and Kaplan 1995; Oswald et al. 2007; Person and Perkel 2005; Reinagel et al. 1999; Smith et al. 2000; Swadlow and Gusev 2001a), we hypothesized that the TRN in an open-loop configuration could have a major impact on information traveling through the TC network.

There is, however, difficulty in measuring information flow through a network of spiking neurons, particularly in networks that must be able to respond to trains of incoming sensory signals at a broad range of rates, such as thalamocortical networks. There exist several generalized methods of estimating dependence between time series (see (Doquire and Verleysen 2012; Silverman 1986; Walters-Williams and Li 2009) for overviews of common estimators). One such class of methods is distance metrics. The Victor-Purpura spike train distance metric (Victor and Purpura 1996) is a binless method of measuring dissimilarity between spike trains. Unlike other distance metrics, this method embeds data in a metric space instead of a vector space, thus avoiding the assumption of spike train addition or scalar multiplication. Another common method is correlation analysis, which includes Pearson correlation and the Spike Timing Tiling Coefficient (STTC) (Cutts and Eglen 2014). Of these methods, STTC was specifically developed for estimating correlation between neural spike trains, and it is not confounded by firing rate, unlike other measures of correlation.

However, a potential problem with traditional correlational analyses is that they are not optimal at estimating nonlinear dependencies (Gencaga et al. 2014), which are often observed in neural networks. This drawback can be addressed by using a dependence measure known as mutual information (MI) (Cellucci et al. 2005; Shannon 1948). MI between two random variables X and Y is mathematically defined as:

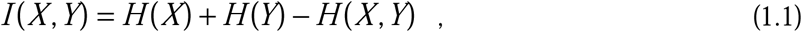

where *H(X)* and *H(Y)* are entropies and *H(X,Y)* is joint entropy. Since the entropy *H* of a random variable *X* = *x*_*1*_,*x*_*2*_,…,*x*_*n*_ is essentially a measure of its unpredictability, and is defined through its probability distribution *p* with 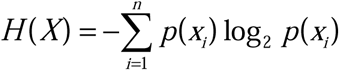, MI estimators can take into account any kind of dependency. Furthermore, an important feature of MI estimators is that they can be used as nonparametric density estimators, as MI is considered to be a nonparametric measure of relevance (Walters-Williams and Li 2009). Nonparametric estimators assume no prior model underlying the data distribution, and consequently require more data than parametric estimators (Worms and Touati 2016). A common example of a nonparametric estimator is the Direct Method (Strong et al. 1998). This estimator calculates MI using the distribution of binary “words” in input and output time series. However, like many estimators the Direct Method uses a uniform partition of the data, which consists of a fixed time window to obtain probability distributions. Uniform partitions are typically ad hoc or chosen through an error-reduction algorithm which increases computational cost (Kjaer et al. 1994; Walters-Williams and Li 2009). By using adaptive partitions (Cellucci et al. 2005; Marek and Tichavsky 2008), convergence of the MI estimate is faster, and the amount of data required is less, making adaptive partition methods generally more computationally efficient than uniform partitions, particularly when data are distributed nonuniformly (Darbellay and Vajda 1999; Marek and Tichavsky 2008; Walters-Williams and Li 2009).

In the case of thalamocortical networks, whose output depends heavily on the rate of synaptic input from both peripheral sensory structures and the TRN (Bartlett and Wang 2007; Kim et al. 1997; Mukherjee and Kaplan 1995; Willis et al. 2015), measurement of the rates of information flow across the thalamus require MI estimators that provide estimates across different spike rates. Therefore, similar to our previous work (Willis et al. 2015), we introduce a modified estimator of MI (<u>A</u>daptive partition using Interspike intervals <u>MI E</u>stimator, or AIMIE), between time series that takes into account these different characteristic time scales. This new estimator, unlike previous estimators, uses adaptive partitions of interspike intervals and spikes densities to handle the disparate firing rates. Previously used estimators are typically limited by poor estimation of nonlinear dependencies, a priori assumptions of distributions, requirements of large amounts of data, and overestimation or underestimation of data distributions. In the current study, the AIMIE method is further examined and compared to these other, more traditional, methods of estimating MI using both simulated and real spike trains. AIMIE is then used to probe the impact of the TRN on the transformation of spike information as signals pass through a simple thalamocortical network model. Using several different criteria, we find that AIMIE outperforms the other metrics and reveals that TRN-mediated inhibition in a thalamocortical model produces a paradoxical recovery of information per spike that is lost during thalamic bursting.

## Methods

### Computational Methods

#### Model Architecture

A Hodgkin-Huxley framework was used to build a three-neuron, open-loop thalamoreticular network, as described previously (Willis et al. 2015). The network consists of single TC, TRN, and L4 cells, modeled as single-compartment models from whole-cell recordings done in our laboratory. Each cell’s membrane potential V was modeled by a first-order differential equation:

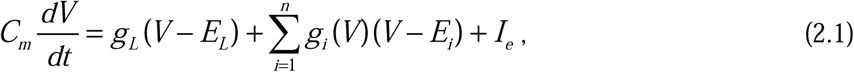

where C_m_ is membrane capacitance, g_L_ is leak conductance, E_L_ is leak reversal potential, I_e_ is current externally applied to the cell, n is the number of channel types, excluding leak channels, g_i_(V) is conductance of ith channel type as a function of the membrane potential, and E_i_ is reversal potential of i^th^ channel type. As described previously (Willis et al. 2015), the model of TC cell includes T-type calcium current, cationic H-current, delayed-rectifying potassium current, and fast sodium current, while the TRN cell model includes all of the aforementioned currents as well as slow-inactivating potassium current (KS current). To detect the maximum number of bursts, the number of bursts in TC output time series was calculated using liberal criteria, defined by (Ramcharan et al. 2000) as, for each burst, at least 50 ms of quiescence followed by at least two spikes with interspike interval(s) of at most 6 ms (but see (Deleuze et al. 2012; Sincich et al. 2007)). The L4 model includes fast sodium current, delayed-rectifying potassium current, and non-inactivating potassium current (M-current). The parameters of these cell models and their currents are found in Table 1. Thalamic afferent, reticulothalamic, and thalamocortical synaptic parameters were derived from the literature (Chen and Regehr 2003; Gentet and Ulrich 2003; Laurent et al. 2002). All inputs to TC and TRN cell models are Poisson-modulated pulse trains, with a single-pulse duration of 0.1 ms.

#### MI Estimator with Adaptive Partition based on Interspike Intervals (AIMIE)

AIMIE is a nonparametric MI estimator (Walters-Williams and Li 2009), which does not assume the distribution of data a priori, similar to the MI estimators mentioned below, and utilizes an adaptive partition (Cellucci et al. 2005) of interspike interval durations and spike densities.

Given two time series of spike times, let X be the time series with the greater number of spikes, and Y be the time series with the lesser number of spikes. Let t_i_^Y^ be the times at which spikes occur in Y, and let Δt_i_^Y^ be the durations (in ms) of an interval between subsequent spikes of Y such that

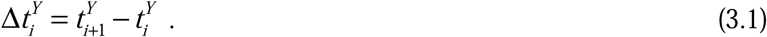

Let n_i_^X^ be the number of spikes of X in the Y interspike interval [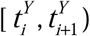) corresponding to Δt_i_^Y^ (Fig 1A), and M be the total number of Y interspike intervals. From Y we constructed a time series of interspike interval durations, Δt_1_^Y^, Δt_2_^Y^, …, Δt ^Y^, and from X we constructed a time series of spike densities, d_1_^X^, d_2_^X^, …, d_M_^X^, corresponding to each interspike interval of Y, where 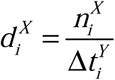.

**Figure 1:**
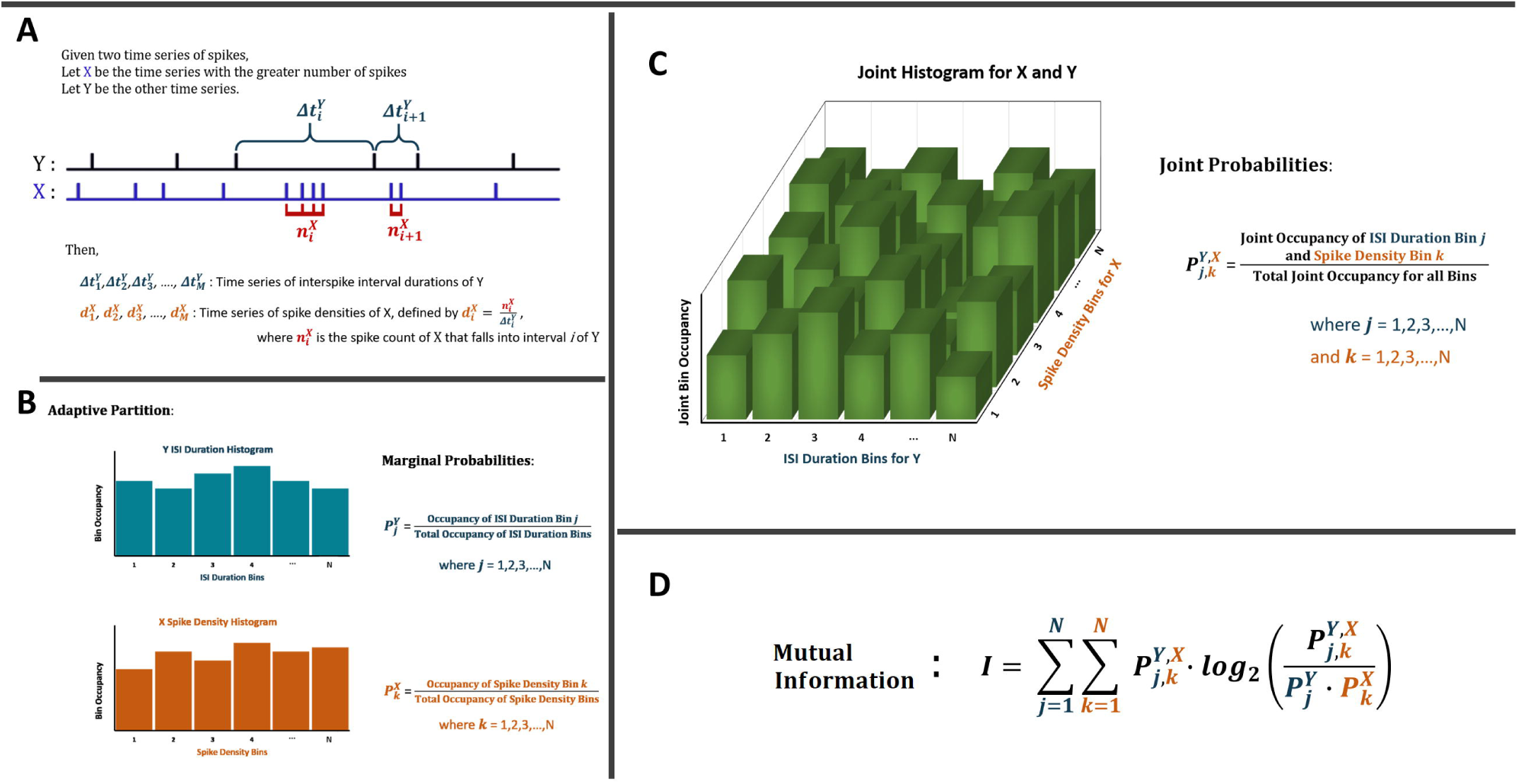
Procedure for estimating MI using AIMIE. A) Given two time series, the one that has a greater number of spikes is designated as X, and the other is designated as Y. A time series of interspike interval durations for Y is constructed, and a time series of spike densities of X corresponding to the interspike intervals of Y. M represents the total number of interspike intervals of Y. B) An adaptive partition is applied to the time series of interspike interval durations of Y and separately to the time series of spike densities of X. Marginal probability for each bin of the adaptive partition is calculated as the occupancy of the bin divided by the sum of occupancies of all bins of that adaptive partition. Note the roughly equal occupancy of the bins, which is due to the adaptive partition. C) A joint histogram is constructed, in which one of the horizontal axes represents the bins of the adaptive partition for Y and the other horizontal axis represents bins of the adaptive partition for X. Joint probability for each combination of bins is calculated as joint occupancy of both bins divided by the sum of all occupancies of the joint histogram. D) Equation for calculating MI, in which the outer sum, from *j*=1 to N, sums over the bins of the adaptive partition of Y and the inner sum, from *k*=1 to N, sums over the bins of the adaptive partition of X.

An adaptive partition was applied to the time series of interspike interval durations of Y and separately to the time series of spike densities of X (Fig 1B). Specifically, for the adaptive partition, the corresponding time series was sorted by smallest to largest values, and each bin was chosen to have a minimum occupancy C_min_ equal to the square root of the total number of interspike intervals of Y, with the total number of bins, for either X or Y, being N. Let C_j_^Y^ be the occupancy of bin j of the adaptive partition used for Y, and let C ^X^ be the occupancy of bin k of the adaptive partition used for X. For each bin, the marginal probabilities, P_j_^Y^ and P ^X^, were calculated as follows (Fig 1B):

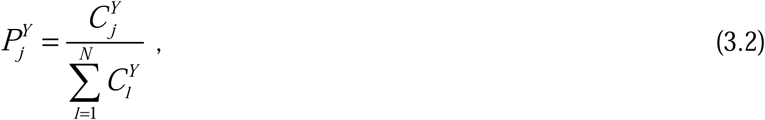

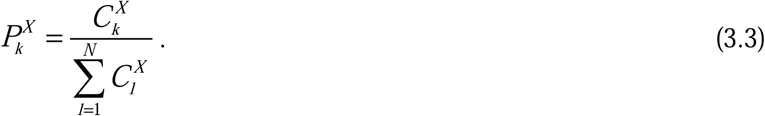

Note that the denominator for each marginal probability is the sum of all occupancies for the corresponding adaptive partition. This particular partition, where each bin is nonempty and has roughly equal occupancy, is related to an adaptive partition (Cellucci et al. 2005).

The joint probability 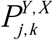 was calculated by constructing a two-dimensional matrix with one horizontal axis corresponding to bins of the adaptive partition for Y and the other horizontal axis corresponding to bins of the adaptive partition for X (Fig 1C). Let 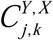 be the joint occupancy of both bin j of the adaptive partition used for Y and of bin k of the adaptive partition used for X. Then 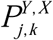 was calculated as follows:

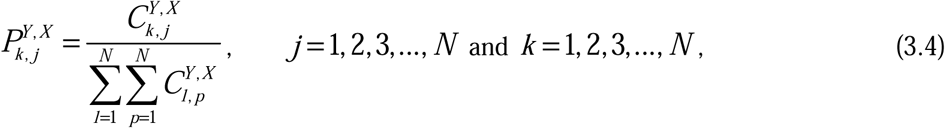

To calculate mutual information, the general formula for MI is used (Fig 1D):

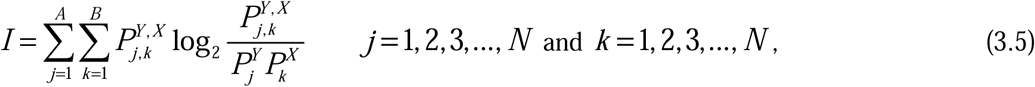

where N, being the total number of bins for either adaptive partition in this case also served as the total number of bins on both axes, such that A = B = N.

#### Direct Method MI estimator (DMIE)

This estimator uses a fixed-width partition of binary input and output time series (Strong et al. 1998). This technique has a precision of Δt, which is the sampling interval that is used to convert input and output time series into binary signals. A time window of length T was used to construct a dictionary of unique binary “words,” each consisting of binary symbols (0’s and 1’s), from the input and output. Marginal probabilities, P_i_^out^ and P_j_^in^, associated with output word i and input word j, were calculated by dividing the total number of occurrences for each word by the total number of occurrences for the corresponding time series. Joint probability, P_i,j_ ^out,in^, was determined by constructing a two-dimensional matrix of output words, corresponding to columns, and input words, corresponding to rows, much like with AIMIE. The occupancy of each two-dimensional bin was then divided by the total occupancy of all two-dimensional bins to give joint probability for the corresponding two-dimensional bin. Finally, MI was calculated using Eq. 3.5, with B as the total number of output words, and A as the total number of input words. We utilized two variations of DMIE: DMIE10, with Δt = 10 ms and T = 100 ms, and DMIE20, with Δt = 20 ms and T = 200 ms.

#### MI estimator with uniform partition of spike counts (three variations: FBWSE, FBNSE, SQRSE)

This is a simple estimator that uses a fixed bin-width partition to determine spike count distributions for input and output. Originally, this method of estimation was used to measure the amount of information transmitted by neuronal responses in the visual system about a set of stimuli (Oram et al. 1999; Oram et al. 2001). Specifically, this estimator can be used to compare information transmitted by total spike count as well as by number of repeating triplets in output spike code. For this method, a uniform partition with time bins of fixed duration, also used for DMIE10, was used to construct a time series of spike counts. Marginal probabilities, P_i_^out^ and P_j_^in^, were determined from histograms that were constructed based on spike counts, with spike bin occupancy defined as the number of time bins containing the corresponding number of spikes. Joint probability, P_i,j_^out,in^, was determined by constructing a two dimensional histogram of input spike bins on one axis and output spike bins on the other axis. MI was calculated using Eq. 3.5, with B as the total number of output spike bins, and A as the total number of input spike bins.

To determine distribution histograms, Oram et. al used the error reduction algorithm of Kjaer et. al (Kjaer et al. 1994; Oram et al. 1999; Oram et al. 2001), which requires training a back-propagation artificial neural network with relatively substantial amounts of data. However, to demonstrate a blind application of this estimator, we defined three variations of uniform partitions that differ only in their construction of time bins. The fixed bin-width spike partition (FBWSE) uses time bins with a fixed duration of 10 ms, regardless of the duration of input or output time series. The fixed bin-number spike partition (FBNSE) uses 500 time bins for both input and output, with bin width being the total duration of the time series divided by number of bins. For the square-root spike partition (SQRSE), we chose the number of time bins to be the square root of the total number of data points for each time series (Cellucci et al. 2005; Mosteller and Tukey 1977).

All simulations and estimators were coded in MATLAB R2012a and R2015a. MI was computed using the algorithms of Celluci et. al (Cellucci et al. 2005). Calculations and simulations were run on multiple machines, including an HP Pavilion machine using a Windows 7 operating system, as well as a Lenovo Ideapad machine with a Windows 10 operating system.

### Electrophysiological Recordings

#### Animals

P20-24 BALB/c mice of both sexes were used for this study. All procedures were approved by the Institutional Animal Care and Use Committee (IACUC, protocol # 16164) at University of Illinois Urbana-Champaign. Animals were housed in animal care facilities at the Beckman Institute for Advanced Science and Technology, approved by the American Association for Accreditation of Laboratory Animal Care (AAALAC).

#### Brain slicing

Mice were initially anesthetized with ketamine (100 mg/kg) and xylazine (3 mg/kg) intraperitoneally and perfused with chilled (4°C) sucrose-based slicing solution ((in mM): 234 sucrose, 11 glucose, 26 NaHCO_3_, 2.5 KCl, 1.25 NaH_2_PO_4_, 10 MgCl_2_, 0.5 CaCl_2_). 300 µm-thick thalamocortical brain slices were obtained, as previously described (Cruikshank et al. 2002; Stebbings et al. 2016), then incubated for 30 minutes in 32°C incubation solution ((in mM): 26 NaHCO_3_, 2.5 KCl, 10 glucose, 126 NaCl, 1.25 NaH_2_PO_4_, 3 MgCl_2_, and 1 CaCl_2_). After incubation, slices were transferred to a perfusion chamber and perfused with artificial cerebrospinal fluid (ACSF) ((in mM): 26 NaHCO_3_, 2.5 KCl, 10 glucose, 126 NaCl, 1.25 NaH_2_PO_4_, 2 MgCl_2_, and 2 CaCl_2_), and bubbled with 95% oxygen/5% carbon dioxide.

#### Electrophysiology

The cell-attached recordings of pairs of neurons located in layer 2/3 or layer 4 of the auditory cortex were performed at room temperature using a visualized slice setup outfitted with infrared-differential interference contrast (IR-DIC) optics. Recording pipettes were pulled from borosilicate glass capillary tubes and had tip resistances of 2–5 MΩ when filled with solution, which contained (in mM): 117 K-gluconate, 13 KCl, 1.0 MgCl_2_, 0.07 CaCl_2_, 0.1 ethyleneglycol-bis (2-aminoethylether)-N,N,N0, N0 -tetra acetic acid, 10.0 4-(2-hydroxyethyl)-1-piperazineethanesulfonic acid, 2.0 Na-ATP, 0.4 Na-GTP, pH 7.3. For data acquisition, a Multiclamp 700B amplifier (Molecular Devices, Sunnyvale, CA, USA) and pClamp software (Molecular Devices, Sunnyvale, CA, USA) were used with a 20-kHz sampling rate. The cell-attached recordings were conducted under after a gigaOhm seal was attained. Once the recording was started, 2 µM of SR-95531 (gabazine, Tocris) was perfused with the ACSF for 30 minutes, after which increasing concentrations of DNQX (Tocris) were added sequentially to the ACSF (approximately 30 minutes/each DNQX concentration) along with 2 µM SR-95531. The software program Clampfit (Molecular Devices, LLC) was used to analyze series of spike times from paired recording data using an event detection algorithm with threshold search.

## Results

The validity of using AIMIE to measure the degree of correlation between spike trains was examined. Four other MI estimators, DMIE, FBWSE, FBNSE, and SQRSE, described in further detail in Methods, are also tested for comparison.

### MI estimator dependence on the length of time series

To determine the number of spikes needed to achieve a stable measurement of MI using AIMIE, simulations of a thalamocortical model were run while systematically varying the simulation time. AIMIE was used to compute the MI between the afferent spike train providing synaptic input to a model TC neuron and the output of the model, measured as spike times in a model layer 4 cortical neuron. A simple thalamocortical network model was used (Fig 2A, see Table 1 for parameters), and the afferent input was a 10 Hz Poisson-modulated spike train, varying only in the total simulation time of the model. This scheme permitted generation of pairs of input and output time series to examine basic properties of AIMIE, such as the dependency of AIMIE on the simulation time, which corresponds to total number of output spikes. Ten trials were performed for each simulation time to calculate standard deviation. Comparing across the total simulation times, MI per output spike approaches an asymptotic value of about 2×10^−4^ bits/spike (Fig 2B). This finding indicates that with increasing amounts of data, AIMIE becomes less dependent on time series length. At about 100 seconds of simulation time (black triangle of Fig 2), which corresponds to an average of 411 output spikes, a decrease is seen in the standard deviation function (Fig 2C) and in magnitude of error bars (Fig 2B inset) when compared to shorter simulation times. Note that for simulation times below 100 seconds there are significant fluctuations in the MI values, however the standard deviation substantially drops at simulation times beyond 100 seconds. Therefore, for most subsequent simulations, simulation times were adjusted such that at least 500 output spikes were generated, and for all tests of AIMIE’s performance, with artificial spike trains that were not generated with the thalamocortical model, a minimum of 500 output spikes was also used.

**Figure 2:**
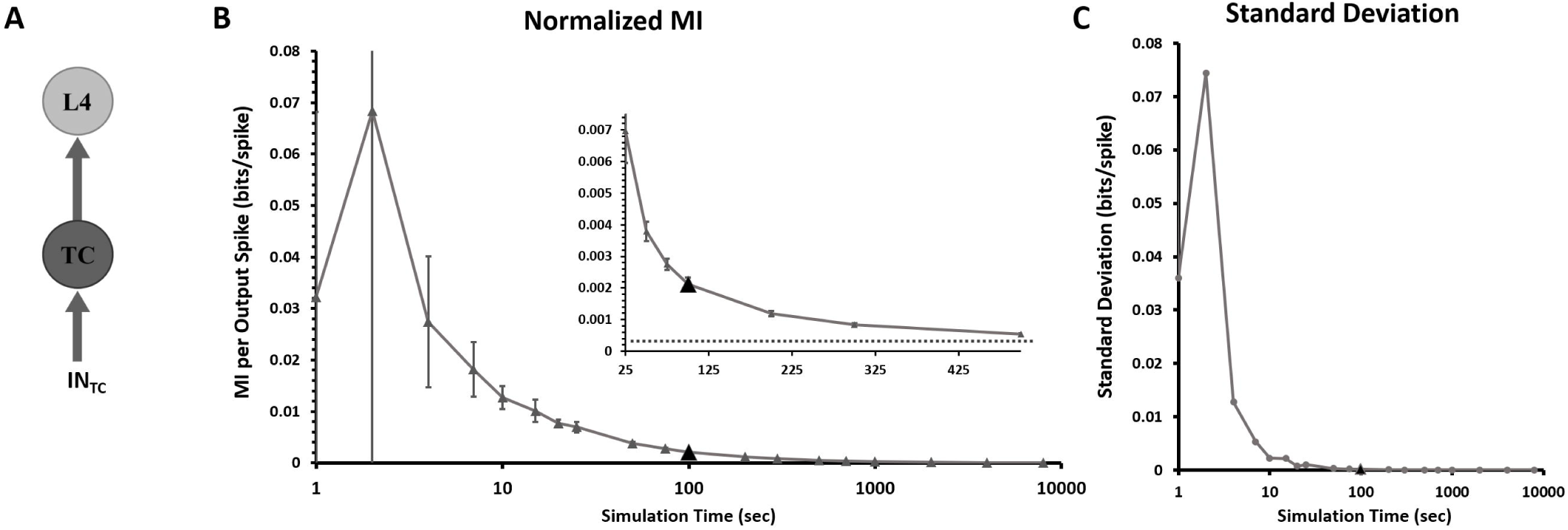
Application of AIMIE to time series generated from the model thalamocortical network with variable simulation times. A) Model architecture of a thalamocortical network containing only TC and L4 neurons, which are modeled using a Hodgkin-Huxley framework. Default model parameters are used for simulations (see Table 1). IN_TC_ corresponds to thalamic afferent inputs, which are generated as 10 Hz Poisson-modulated pulse trains. B) MI per output spike provided by AIMIE when it is applied to time series pairs with different simulation times. As simulation time increases, the number of output spikes increases as well, and AIMIE’s MI per output spike trends towards a horizontal asymptote of about 2×10^−4^ bits/spike. The black triangle marks a simulation time of 100 seconds, which provides an average of about 411 output spikes. Inset shows decreased fluctuations in MI per output spike on an expanded scale, particularly for simulation times above 100 seconds. 10 trials were used for each simulation time, to generate error bars and standard deviation. C) Standard deviation of AIMIE’s estimates of MI per output spike from Fig 2A for a range of simulation times. Note that standard deviation is significantly smaller at around 100 seconds of simulation time (black triangle) than at shorter simulation times.

### MI estimator independence of interspike interval length variation

To determine whether any of the MI estimators are sensitive to the average spiking rate, we created artificial input and output spike trains and computed MI between these trains using multiple different MI estimators. The input time series were randomly generated using a uniform distribution, each with 4000 spikes in total and interspike intervals ranging from 5 ms to 50 ms. In this case, for the output time series, a single output spike followed each input spike and arrived before the next input spike. Under the assumption here that the optimal estimator for MI should be insensitive to stretch (and therefore average spiking rate), the input and output spike trains were then temporally “stretched” to change their average spiking rate without changing the relationship between the two trains (Fig 3A). In total, ten trials were performed for each average spiking rate. When tested alongside the four other estimators (Fig 3B), the estimators that produced a change in MI beyond 10% of the baseline MI were DMIE10, DMIE20, and FBWIE, all of which use a fixed bin width partition, indicating that DMIE and FBWIE are sensitive to time scaling, while FBNIE, SQRSE, and AIMIE are not. Therefore, FBNIE, SQRSE and AIMIE were compared in subsequent examinations of MI estimator performance.

**Figure 3:**
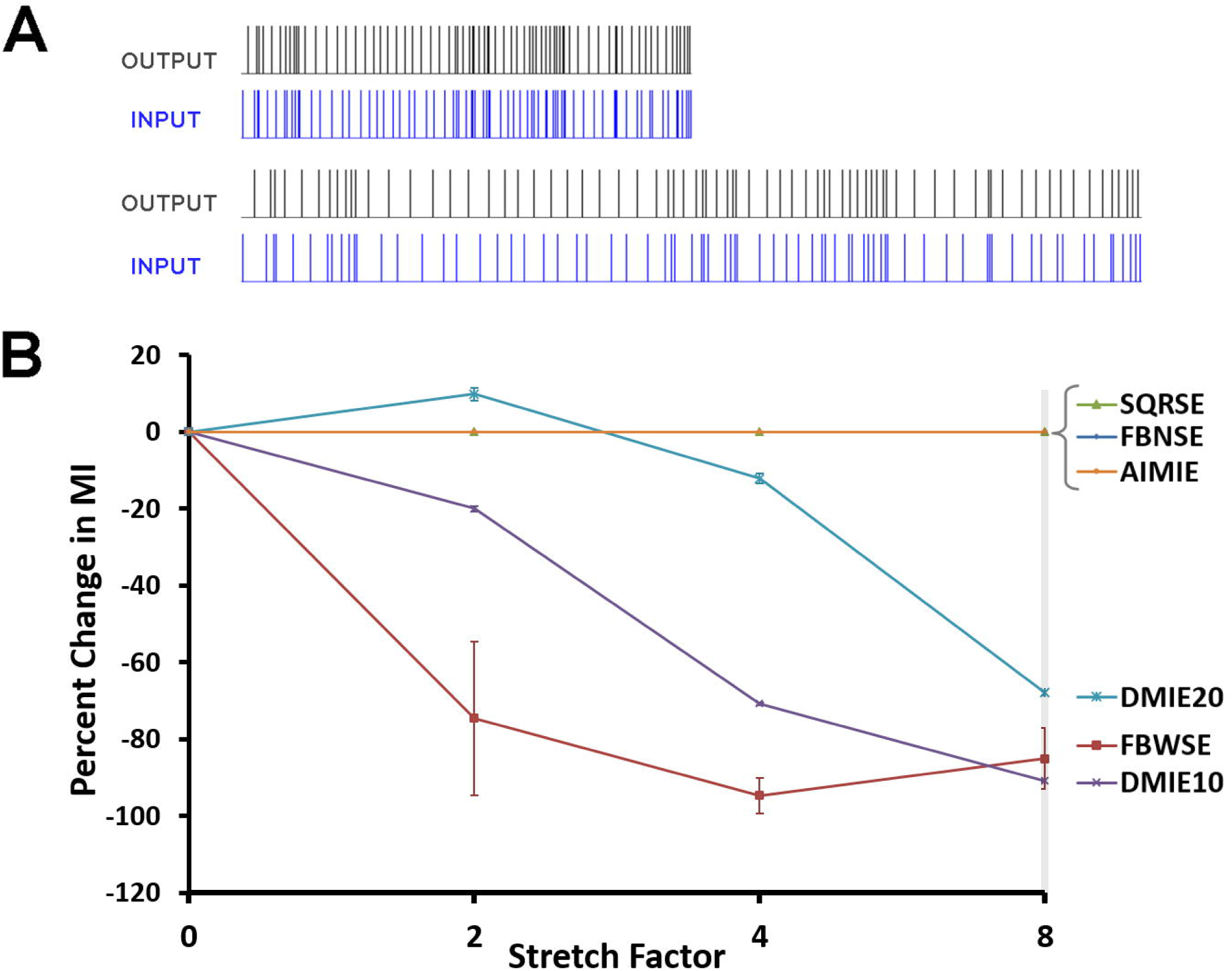
Effect of scaling of interspike intervals on MI for different estimators. A) Spike time plots demonstrating scaling of interspike intervals for input and output series. In this case, the second input and output pair is generated by scaling all interspike intervals of the first pair by a factor of 2. B) Percent change in MI demonstrated by estimators AIMIE, DMIE, FBWSE, FBNSE, and SQRSE when they are applied to input and output time series with interspike intervals scaled by factors of 2, 4, and 8. 10 trials were used for each data point and to generate standard deviation for error bars.

### MI estimator performance with artificial spike trains

In the next test of MI estimator performance, it was assumed that a high dependence between input and output data would be indicated by each input spike eliciting a similar spiking event in the output. Deviation from this idealized response would be considered a loss of information. To test the performance of the MI estimators, four sets of input and output time series were generated, with progressively decreasing degree of correlation in the first three sets (Fig 4A). The input time series were randomly generated, each with 4000 spikes in total and interspike intervals ranging from 5 ms to 50 ms using a uniform distribution. For the first set of input and output time series, designated as Type 1, the response is ideal, meaning that each input spike has 100% chance of eliciting a single output spike. For the Type 2 pair, there is a 50% response, where each input spike has a 50% chance of generating a single output spike. This adjustment roughly decreases the number of output spikes in Type 2 by half when compared to Type 1, and as such, should constitute a decrease in MI. Similarly, the Type 3 pair features a 25% response. The pair of time series in Type 4 is constructed similarly to Type 3 time series, with the 4000 inputs randomly generated; each input has a 25% chance of generating a response event in the output, which, unlike for Type 3, is a random number between 1 to 10 spikes. Ten trials were performed for each application of an MI estimator to a different type of time series pair. Our expectation was that an ideal MI estimator would show the highest MI for Type 1 stimuli and progressively lower for all of the rest.

**Figure 4:**
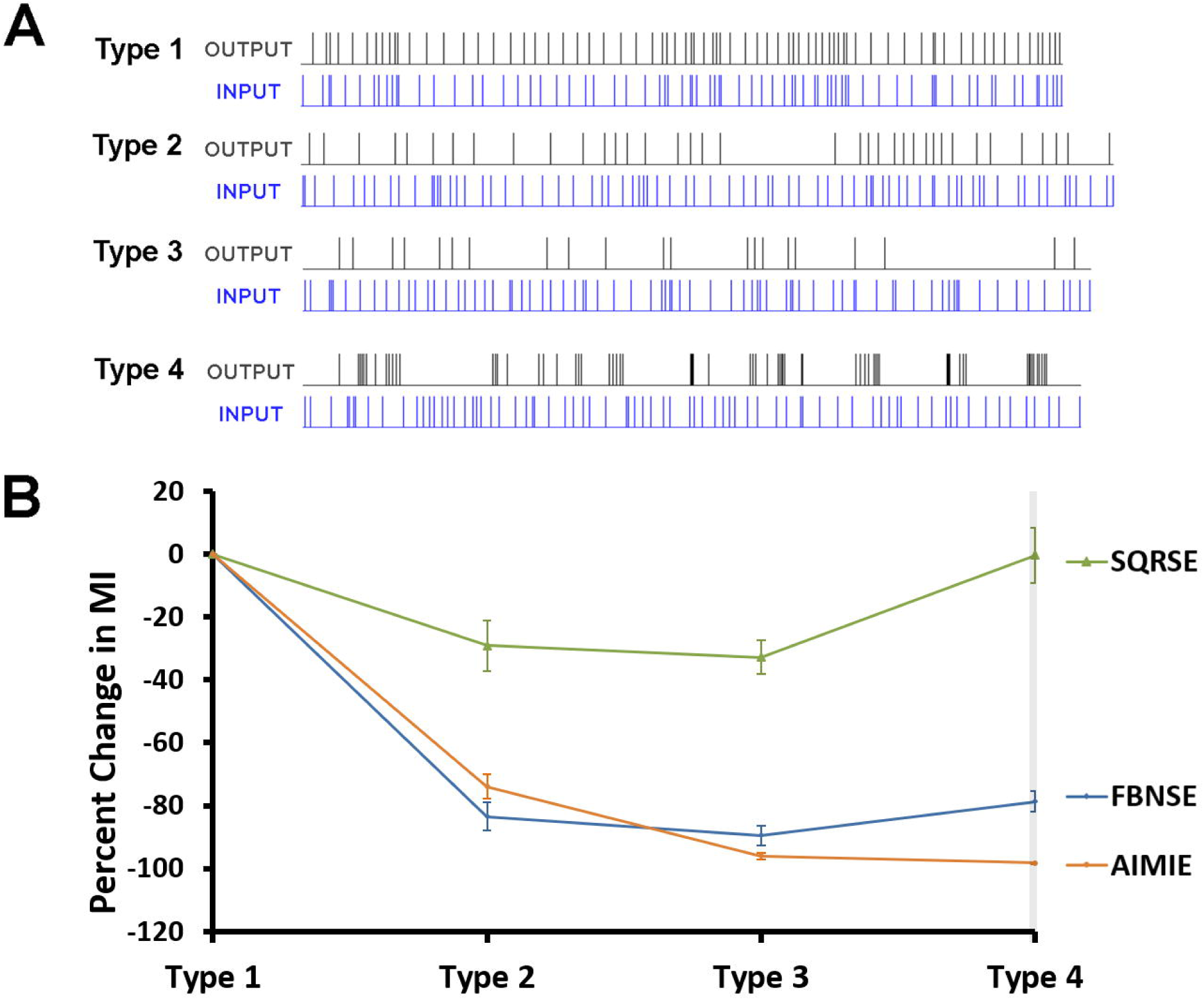
Application of different estimators to hypothetical input and output series of variable dependence. A) Spike time plots of hypothetical constructed time series of Types 1, 2, 3, and 4, where Type 1 is an ideal (100%) response type, Type 2 is a 50% response type, Type 3 is a 25% response type, and Type 4 is a 25% burst response type. B) Percent change in MI demonstrated by estimators AIMIE, FBNSE, and SQRSE when they are applied to input and output time series of Types 1, 2, 3, and 4. As before, 10 trials were used for each data point and to generate standard deviation for error bars.

For this comparison, only the MI estimators that were shown to be insensitive to time scaling (FBNIE, SQRSE, and AIMIE) were tested. When AIMIE was applied to these four pairs of constructed time series, it was observed that the MI values corresponded to the drops in degree of correlation across Types 1 through 3 (Fig 4B). For Type 4, the MI value was very close to that of Type 3, within 2%. FBNSE and SQRSE, neither of which uses an adaptive partition, demonstrated a significant drop in MI across Type 1 to Type 2 and a slight drop in MI from Type 2 to Type 3. Lastly, for Type 4, when compared to Type 3, both FBNSE and SQRSE showed an increase in MI. Note that the only estimator whose MI value for Type 4 was less than that of Type 3 is AIMIE. Substantial fluctuation in MI values, resulting in significant error values, may indicate an insufficient amount of data, particularly for SQRSE, though as suggested by our time series length test (Fig 2). This amount of data would be enough for AIMIE to yield reliable results, as there are 4000 input spikes and around 1000 or more output spikes and for each time series pair type. Note that the degree of variance for AIMIE’s estimates is noticeably smaller than those of the other estimators’ values. These data suggest that AIMIE is more sensitive than other methods to manipulations of spike trains that are expected to diminish the MI between them.

### MI estimator performance with paired electrophysiological recordings

In this test, MI estimators were applied to spike trains of paired recordings of spontaneous activity from the auditory cortex in a slice preparation, under different concentrations of bath-applied DNQX, which is an AMPA receptor antagonist that blocks excitatory synaptic transmission. At higher concentrations of DNQX, we expect that MI per spike between responses of the two neurons of a paired recording would drop, due to diminished synchrony between spontaneous action potentials of the paired neurons. Specifically, AIMIE and FBNSE were tested, as these two estimators performed well with artificially constructed inputs and outputs described above. Recordings from a total of five pairs of neurons in layer 2/3 or layer 4 from mouse brain slices containing the auditory cortex were used. For each pair of recorded neurons, the neuron that produced the greatest number of spikes at the highest DNQX concentration was used for normalization of MI, specifically by dividing the MI value of each trial by the number of spikes produced by this neuron for that trial. For each paired recording sequence under different DNQX concentrations, the change in normalized MI was calculated by dividing the MI per spike at each DNQX concentration by the MI per spike at DNQX concentration of 0 µM. With increasing DNQX concentrations, AIMIE demonstrated the expected drop in normalized MI (Fig 5A), while FBNSE did not (Fig 5B), indicating that AIMIE performs well with electrophysiological recordings of variable synchrony.

**Figure 5:**
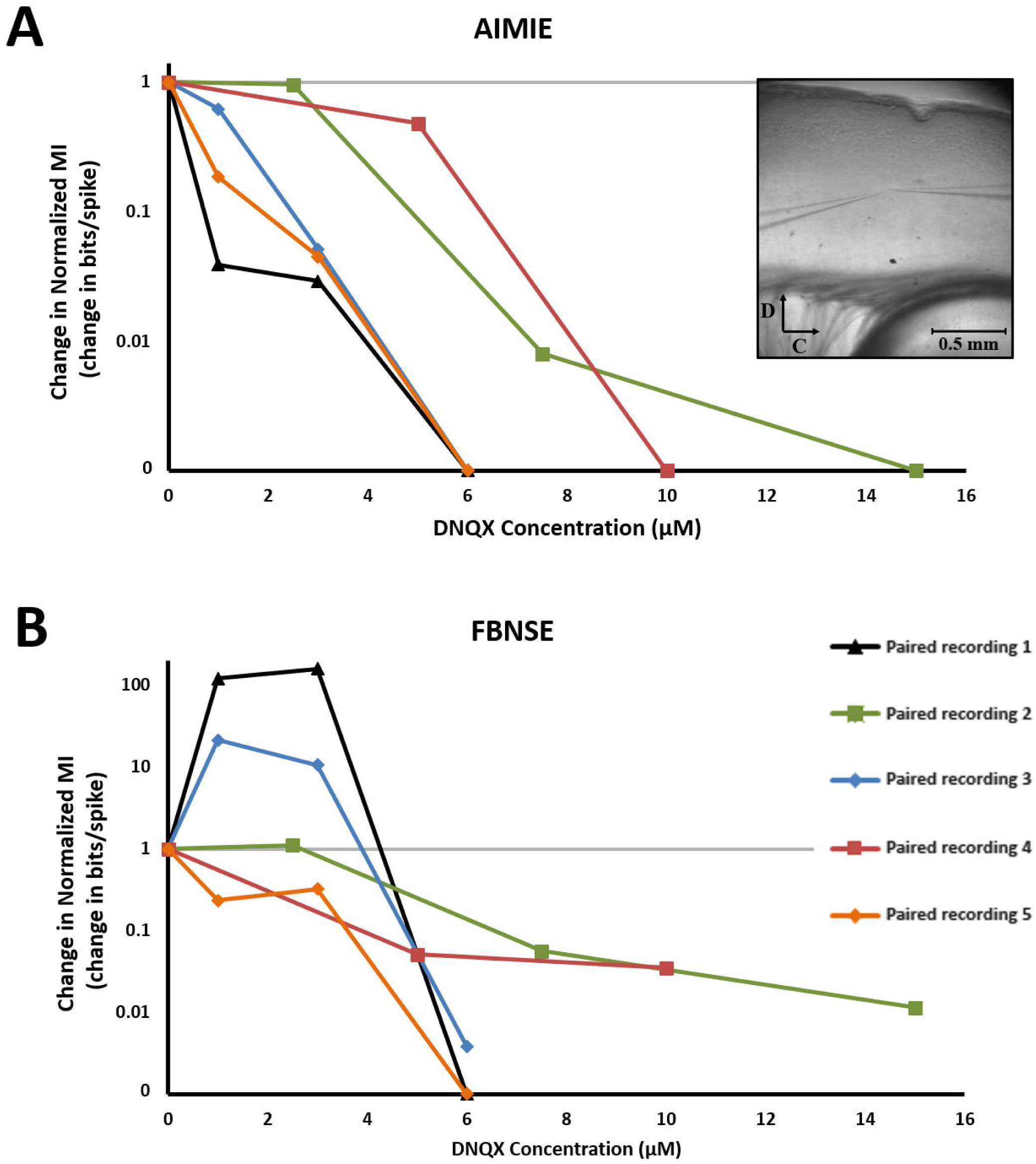
Application of different estimators to dual-recorded spike time series of variable synchrony. Recordings from a total of five neuron pairs are used. For both estimators, for each paired recording sequence under different DNQX concentrations, the change in normalized MI is calculated as MI per spike at each DNQX concentration divided by the MI per spike at DNQX concentration of 0 µM. A) Change in normalized MI demonstrated by AIMIE across different concentrations of DNQX. The inset is an image of electrode placement for a paired recording in auditory cortex of mouse brain slice, with C denoting caudal and D denoting dorsal orientation. The legend for paired recordings is located below in Fig 5B. B) Change in normalized MI demonstrated by FBNSE across different concentrations of DNQX. The legend corresponds to both Fig 5A and Fig 5B.

### Varying T-current conductance in open-loop thalamoreticular network

Applying AIMIE to a simple open-loop thalamocortical network model (Fig 6A), the effect of varying stimulation parameters and the T-current conductance of the TC cell on the modification of an ascending signal through the thalamus was explored. Poisson-modulated synaptic input to a model TC cell (henceforth “thalamic afferents”) was varied from 0.5 Hz to 200 Hz on a logarithmic scale, while Poisson-modulated synaptic input to a TRN neuron was similarly varied from 0 Hz to 200 Hz. As before (Willis et al. 2015), the applied spike trains in thalamic afferents were independent of the applied spike trains in afferents to the TRN cell. For each combination of TC and TRN stimulation rates, the MI between the thalamic afferents and output, either at the TC or L4 neuron, was computed. This MI, normalized by the number of output spikes, was then averaged over all combinations of TC and TRN stimulation rates to produce average MI per output spike. The average MI per output spike was computed for each T-current conductance, ranging from 0 to 100 nS (Fig 6B). Also, for each value of T-current conductance, ranging from 10 to 80 nS, a heat map plot of normalized MI values with TRN stimulation rates on the x-axis and TC stimulation rates on the y-axis was constructed (Fig 7A for normalized MI between thalamic afferents and L4 output, and Fig 7B for normalized MI between thalamic afferents and TC output). 10 trials were performed for each data point of T-current conductance.

**Figure 6:**
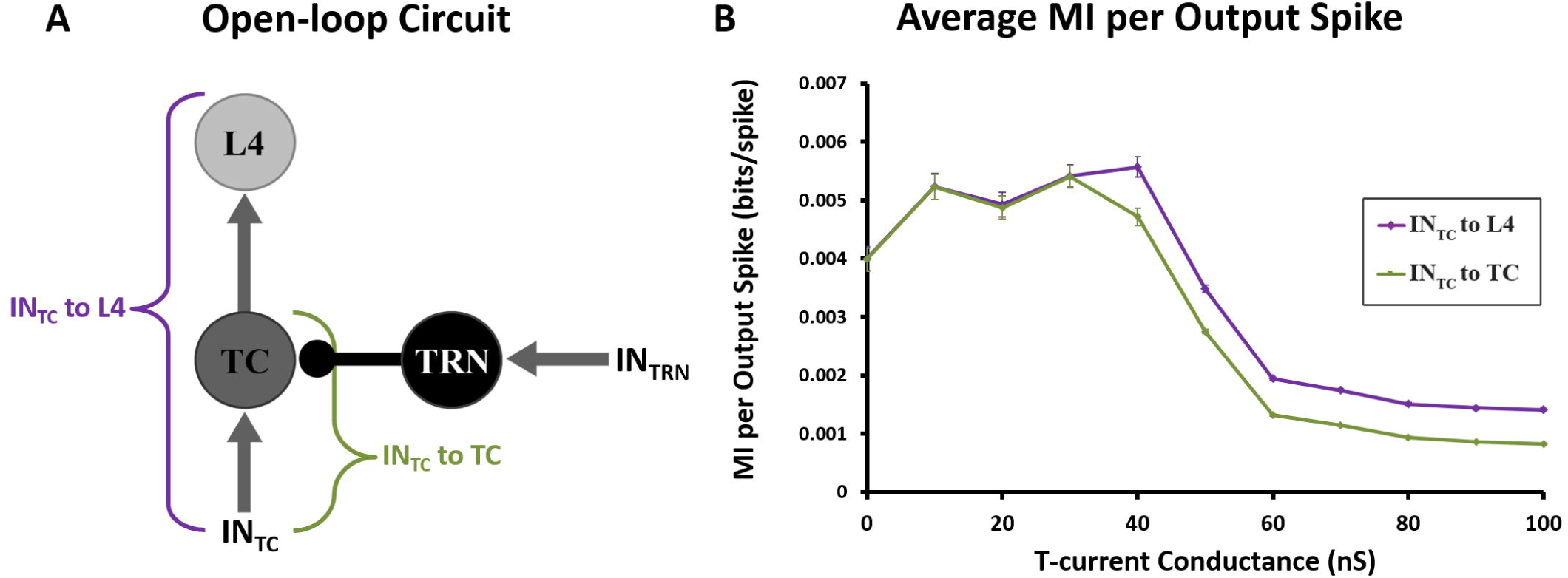
MI analysis of inputs and outputs in an open-loop thalamocortical network model at variable T-current conductances. A) Model architecture of the open-loop thalamocortical network. Arrows represent excitatory inputs, while the TRN to TC projection represents inhibitory (GABAergic) input. All neurons are modeled using a Hodgkin-Huxley framework. IN_TRN_ corresponds to input to the TRN, which ranges from 0 Hz to 200 Hz, while IN_TC_ corresponds to thalamic afferents, which ranges from 0.5 Hz to 200 Hz. Both IN_TRN_ and IN_TC_ are generated as Poisson-modulated pulse trains. Note that the green bracket symbolizes information transfer from thalamic afferents to TC, and the purple bracket symbolizes information transfer from thalamic afferents to L4. B) Average MI transmitted per output spike between thalamic afferents and TC and between thalamic afferents and L4, at a range of TC T-current conductances. There is a peak in MI per output spike at a TC T-current conductance of about 40 nS, which may indicate maximum potentiation of ascending input in open-loop thalamocortical network. Again, 10 trials were used for each data point and generation of standard deviation for error bars.

**Figure 7:**
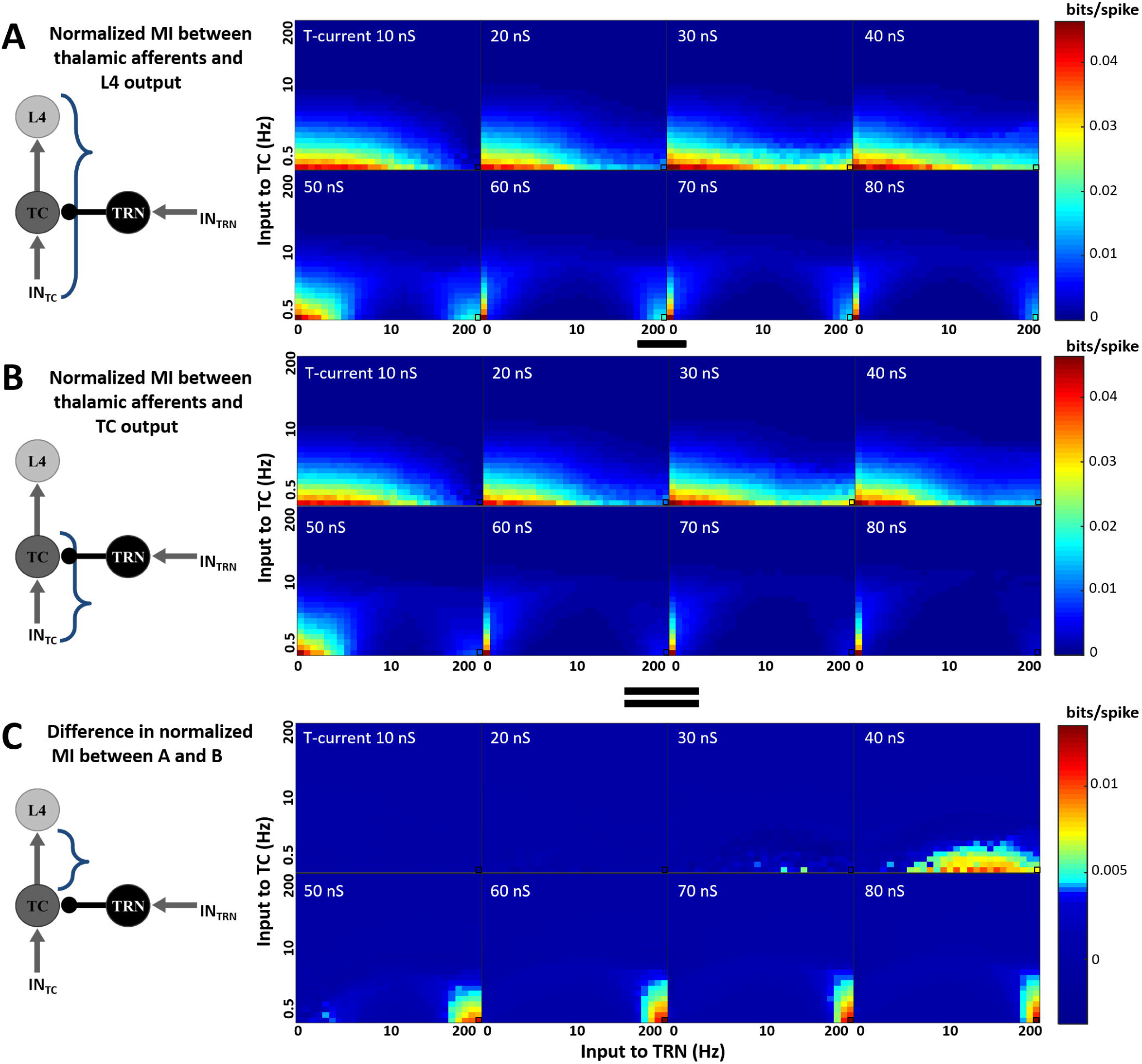
Heat map plots of normalized MI in open-loop thalamocortical network model for variable T-current conductances. For each TC T-current conductance, an MI plot is generated using a range of combinations of TC and TRN stimulation rates and then normalized by the number of output spikes for each rate combination. Normalized MI plots are averaged over 10 trials. The rate combination of 207 Hz stimulation of TRN and 0.5 Hz stimulation of TC (see Fig 8) is marked in a black box on each plot. Note that the horizontal black line between panels A and B denotes a minus sign, while the pair of horizontal black lines between panels B and C denotes an equals sign. A) Plots of average normalized MI (bits per output spike) between thalamic afferents and L4 output for T-current conductances of 10 nS to 80 nS. B) Plots of average normalized MI between thalamic afferents and TC output for T-current conductances of 10 nS to 80 nS. C) Plots of average normalized MI between thalamic afferents and L4 output (Fig 7A) minus average normalized MI between thalamic afferents and TC output (Fig 7B) for T-current conductances of 10 nS to 80 nS. These difference plots show a recovery of information per spike at low thalamic afferent rates and high TRN stimulation rates.

The average MI per output spike between the thalamic afferents and L4 is on average slightly greater than between the thalamic afferents and the TC cell (Fig 6B). Note that at certain combinations of TRN and TC stimulation rates, specifically at high TRN stimulation frequencies and low TC stimulation frequencies, MI per output spike between the thalamic afferents and L4 output paradoxically becomes greater than MI per output spike between the thalamic afferents and TC output, as seen in the normalized MI difference plots of Fig 7C. For example, for TC T-current conductances above 20 nS and thalamic afferent input at 0.5 Hz and to the TRN neuron at 207 Hz, MI per spike across the whole network (from thalamic afferent to L4; purple line, Fig 8A) paradoxically exceeds that seen during the first stage of the network (from thalamic afferent to TC; green line, Fig 8A). In addition, this paradoxical behavior is seen as the degree of bursting of the TC neuron increases (blue line, Fig 8A). This relationship is quantified in Fig 8B. Here, the number of bursts in the TC neuron is compared against this paradoxical increase in MI across the whole network, and a strong positive correlation is observed (Fig 8B, (Pearson’s r = 0.843, p < 0.001)). These data suggest that the paradoxical increase in MI per spike is related to the underlying bursting behavior of the TC cell.

**Figure 8:**
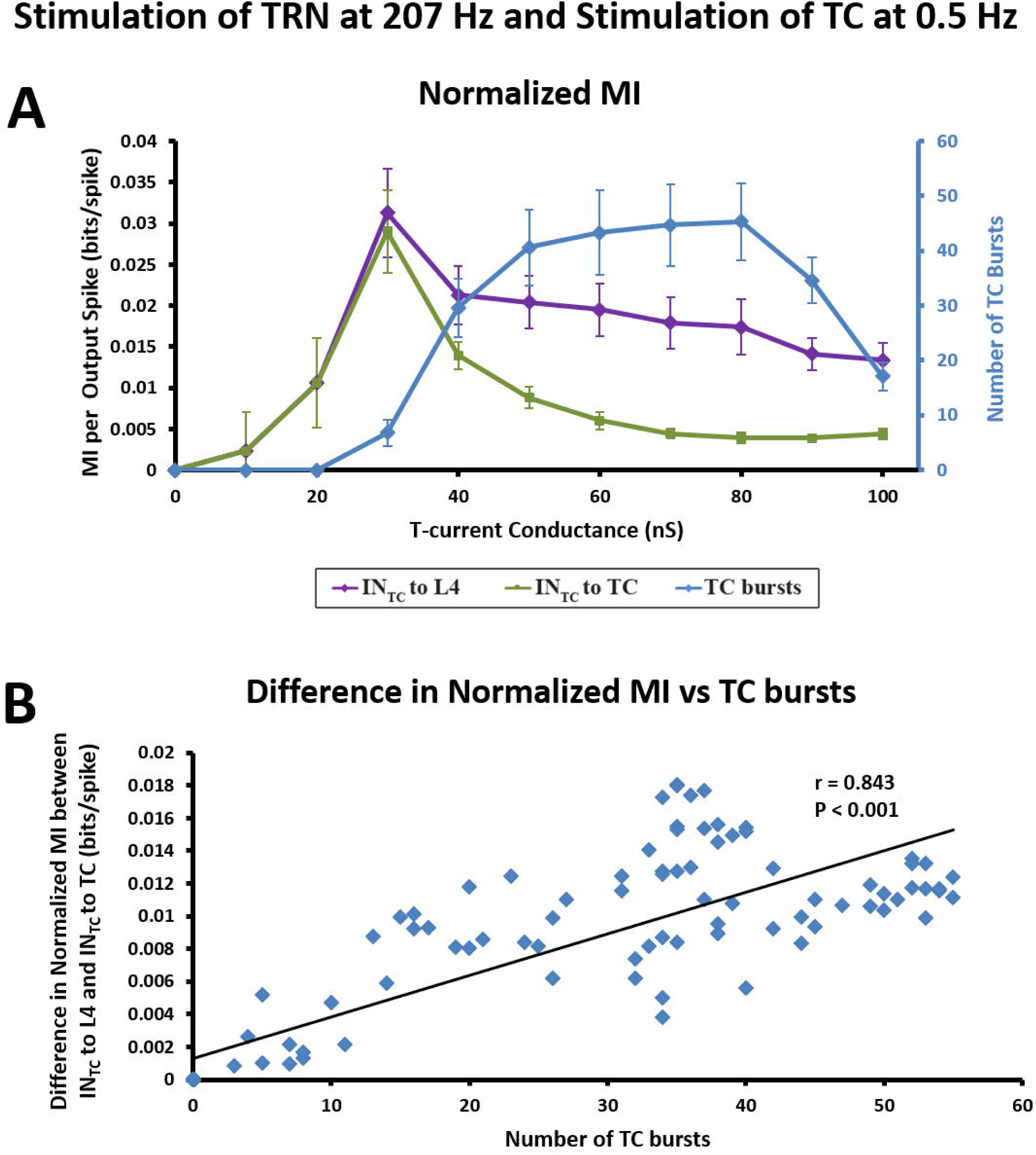
Stimulation of TRN at 207 Hz and TC at 0.5 Hz in open-loop thalamocortical network model. This rate combination is marked as a black box on the normalized MI plots of Fig 7. IN_TC_ refers to thalamic afferents, as shown in Fig 7. A) MI per output spike between thalamic afferents and TC output and thalamic afferents and L4 output at a range of T-current conductances from 0 nS to 100 nS. The number of bursts observed at the TC is also shown alongside on the right vertical axis. 10 trials were used for each data point and for generation of standard deviation for error bars. B) Difference of MI per spike of TC input to TC output and MI per spike of TC input and L4 output against the number of bursts produced by TC. The trendline is shown alongside a Pearson coefficient value of r = 0.843 (p < 0.001).

## Discussion

### Summary of findings

In the current study, the properties of a novel estimator of MI, the <u>A</u>daptive partition using Interspike intervals <u>MI E</u>stimator (AIMIE), were examined, with its performance compared to other, more established methods of measuring MI between spike trains. The AIMIE method of computing MI was found to be insensitive to overall time compression or expansion of the spike trains, suggesting that it may be used effectively at both high and low spike rates. AIMIE was also found to be sensitive to manual manipulation of the relationship between simulated input and output spike trains. That is, manual degradation in the relationships between two spike trains also lowered the MI, computed with AIMIE, but not with other methods. Further, in paired neuronal recordings, the decrease in MI with increasing concentration of DNQX was consistent with DNQX’s effect of desynchronizing the responses of the recorded neuron pairs. Finally, when AIMIE was used to compute the MI between input (to a model TC cell) and output (from a model L4 neuron) in a thalamocortical model, it was found that MI was maximum at T-current amplitudes corresponding to those seen physiologically, and that the apparent degradation in MI caused by bursting could be recovered at the thalamocortical synapse. These data suggest that a new method for estimating MI between spike trains, AIMIE, is now available to investigators and that the method has the advantage of being able to easily compensate for wide changes in the average spike rate of either the input or output spike train.

### Methodological considerations

Since the thalamocortical network demonstrates short latencies in L4 neurons in response to afferent input provided to TC neurons (< 10 ms, corresponding to 2 synapses between input and output, (Llano et al. 2014), no effort was made to shift the output time series relative to the input time series. However, in other networks, particularly in large-scale networks containing many more synapses, where response latencies are longer, a constant time delay may need to be compensated for in the analysis. In addition, most MI estimators require a minimum amount of data for reliable results (Paninski 2003). As observed in Fig 2B and C, under the specified simulation parameters, approximately 400-500 output spikes, corresponding to 100 seconds simulation time, are needed for AIMIE to provide consistent MI values, suggesting that for spike rates of approximately 10 spikes/second, approximately 40-50 seconds of data are required to compute MI using AIMIE. Parametric density estimators, such as maximum likelihood estimation, require relatively little data for convergence (Marek and Tichavsky 2008), while MI estimators with uniform partition, especially DMIE, usually require larger amounts of data in comparison to MI estimators with adaptive partition (Borst and Theunissen 1999; Walters-Williams and Li 2009).

### MI as an estimate of dependence

Multiple metrics have been used previously to compute the degree of similarity between spike trains. MI is a quantity that measures general dependence between two random variables, and as such, it is related to correlation functions which traditionally measure linear dependence (Li 1990). STTC, for example, uses a fixed time bin around each input spike to calculate correlation between input and output spike trains (Cutts and Eglen 2014). This approach is reminiscent of a uniform partition method, which is inadequate for analyzing MI in a TC network because of the wide range of input rates processed by TC neurons, discussed below. Related to the correlation measures are distance metrics, including the Victor-Purpura spike train distance metric (Victor and Purpura 1996). This metric is commonly used in input and output spike train analysis, though there is evidence that this may not be an optimal metric for evaluating temporal information (Gai and Carney 2008). Specifically, the Victor-Purpura spike train distance metric measures information carried primarily by absolute spike times, and it does not appear to effectively account for other temporal features in spike time series, such as synchronization to frequency of sensory input in a neuron’s response. This insensitivity to certain temporal features could potentially complicate its use in analysis of the temporally sensitive thalamocortical network.

Furthermore, traditional correlation analysis, such as Pearson correlation, has been shown to be optimal only in the case of linear dependencies between variables, while MI can be applied to random variables that exhibit linear as well as nonlinear dependencies (Gencaga et al. 2014), which are often demonstrated in neural networks (Schöner and Kelso 1988). Comparisons of marginal entropy, meaning the average amount of information provided by, in this case, a single spike time series, have also been used to measure the relative information content of time series in the analysis of bursting and tonic firing in the thalamus (Reinagel et al. 1999). However, only marginal entropies are compared in this earlier analysis, and there is no calculation of joint entropy between two time series, as there is in MI estimation. A consequence of not calculating joint entropy is that the relative timing between input and output spikes is lost.

Many previously employed MI estimators use uniform partitions of data to estimate density distributions (e.g., DMIE, FBNSE, FBWSE, and SQRSE). While computationally efficient, these uniform partitions can produce many empty bins (Gencaga et al. 2014). Therefore, for these estimators, different choices of bin construction for a nonadaptive partition can produce substantially different MI values. Estimators with a fixed bin width that do not depend on the number of data points, FBWSE and DMIE, show a change in MI when applied to time-scaled time series (Fig 3), unless bin widths of the partition are scaled appropriately. Furthermore, estimators with a fixed bin width that do depend on the number of data points, FBNSE and SQRSE, did not perform as well as AIMIE with certain pairs of time series of variable degrees of correlation and synchrony (Figs 4 and 5). These findings regarding estimators that use fixed bin widths indicate that in a network where spike timing can be changed significantly by varying input rate, an MI estimate of each time series pair would require a readjustment of the uniform partition. The use of multiple partition schemes could cause substantial uncertainty in comparisons across samples, since every combination of TRN and TC stimulation rates would require its own parameters for the uniform partition. Therefore, for the thalamocortical network, MI estimators with uniform partitions would not be an optimal choice, leaving us to favor an adaptive partition instead. In addition, unlike estimators that use a uniform partition, AIMIE’s adaptive partition also causes estimated MI to converge faster with sample size (Walters-Williams and Li 2009), and allows investigators to work with time processes that have incomparable time-scales (Darbellay and Vajda 1999).

### T-current conductance, bursting, and MI

Systematic variation of the TC cell’s T-current conductance in the thalamocortical model revealed that there is a specific range of maximal conductance values near 40 nS that produces a maximum average MI per output spike between thalamic afferent input and output at L4 (Fig 6B). This finding suggests that the TRN is able to induce the most potentiation of the ascending TC input at this peak TC T-current conductance value. Note that this value is close to the physiological TC T-current conductance of approximately 45 nS (Willis et al. 2015), and is within the range used in previous modeling studies (Deleuze et al. 2012; Destexhe et al. 1993; Pospischil et al. 2008; Wang 1994). It has been shown that some hyperpolarization-activated cation currents can be modulated by factors such as intracellular pH and specific neurotransmitters, including norepinephrine and serotonin (Munsch and Pape 1999; Pape and McCormick 1989). If TC T-current is similarly modulated (for example see (Joksovic et al. 2010)), then the current study suggests that this modulation could alter the amount of MI between input and output as well. Thus, findings from the current study may indicate the possibility that modulation of TC T-current is one way in which the thalamocortical network can modulate sensory thalamocortical information flow.

Previous studies have suggested that bursting in thalamic relay neurons degrades ascending sensory signals, and that tonic firing modes are more likely to produce a high-fidelity representation of ascending information en route to the cortex (Castro-Alamancos 2002; McCarley et al. 1983; McCormick and Feeser 1990). However, it has also been shown that thalamic bursts may carry more information, can enhance detection of specific temporal sequences, and are higher in synaptic efficacy than single thalamic spikes (Lesica and Stanley 2004; Lesica et al. 2006; Reinagel et al. 1999; Swadlow and Gusev 2001a). The current study suggests that in certain cases, particularly with high-frequency input to TRN and low frequency input to TC (Fig 8A), bursting is responsible for loss of incoming sensory information at the level of the TC (Fig 8B), as it likely obscures the ascending sensory signal. However, this information loss may be compensated in the L4 neuron due to the filter properties of the TC synapse, which produce single spikes in response to the initial spikes of a burst (Boudreau and Ferster 2005; Chung et al. 2002; Gil et al. 1999; Krahe and Gabbiani 2004; Swadlow and Gusev 2001b). Thus, information per spike that is lost at the level of the thalamus due to bursting may be partially recovered at the level of the cortex; this is a hypothesis that can be tested physiologically in subsequent studies.

## Acknowledgements

The authors thank Drs. Rama Ratnam and Jeffrey Brown (University of Illinois) for their reviews of this manuscript prior to submission. This work was supported by DC013073 and DC014765 to DAL.

