## Supplementary Materials for "A novel mutual information estimator to measure spike train correlations in a model thalamocortical network"

| **TC Cell** | |
| --- | --- |
| Leak conductance | 3.263 nS |
| Leak equilibrium potential | -60.03 mV |
| Fast sodium conductance | 1500 nS |
| Sodium equilibrium potential | 50 mV |
| Potassium conductance | 520 nS |
| Potassium equilibrium potential | -100 mV |
| T-current conductance | 45 nS |
| Calcium equilibrium potential | 120 mV |
| H-current conductance | 0.608 nS |
| H-current equilibrium potential | -33 mV |
| **L4 Cell** | |
| Leak conductance | 4.8128 nS |
| Leak equilibrium potential | -60.2354 mV |
| Fast sodium conductance | 3000 nS |
| Sodium equilibrium potential | 50 mV |
| Potassium conductance | 140 nS |
| Potassium equilibrium potential | -90 mV |
| M-current conductance | 1.5 nS |
| **TRN Cell** | |
| Leak conductance | 3.7928 nS |
| Leak equilibrium potential | -57 mV |
| Fast sodium conductance | 3000 nS |
| Sodium equilibrium potential | 50 mV |
| Potassium conductance | 400 nS |
| Potassium equilibrium potential | -100 nS |
| T-current amplitude | 21 nS |
| H-current amplitude | 0.0192 nS |
| H-current equilibrium potential | -33 mV |
| KS-current conductance | 3.5 nS |
| KS current τ | 200 ms |

**Table 1: Thalamocortical Network Model Cellular Parameters.**
